# Biomorphogenic feedbacks and the spatial organisation of a dominant grass steer dune development

**DOI:** 10.1101/2021.08.02.454713

**Authors:** Dries Bonte, Femke Batsleer, Sam Provoost, Valérie Reijers, Martijn L. Vandegehuchte, Ruben Van De Walle, Sebastian Dan, Hans Matheve, Pieter Rauwoens, Glenn Strypsteen, Tomohiro Suzuki, Toon Verwaest, Jasmijn Hillaert

## Abstract

Nature-based solutions to mitigate the impact of future climate change depend on restoring biological diversity and natural processes. Coastal foredunes represent the most important natural flood barriers along coastlines worldwide, but their area has been squeezed dramatically because of a continuing urbanisation of coastlines, especially in Europe. Dune development is steered by the development of vegetation in interaction with sand fluxes from the beach. Marram grass (*Calamagrostis arenaria*, formerly *Ammophila arenaria*) is the main dune building species along most European coasts, but also in other continents where the species was introduced. Engineering of coastal dunes, for instance by building dunes in front of dikes, needs to be based on a solid understanding of the species’ interactions with the environment. Only quantitative approaches enable the further development of mechanistic models and coastal management strategies that encapsulate these biomorphogenic interactions. We here provide a quantitative review of the main biotic and physical interactions that affect marram grass performance, their interactions with sand fluxes and how they eventually shape dune development. Our review highlights that the species’ spatial organisation is central to dune development. We further demonstrate this importance by means of remote sensing and a mechanistic model and provide an outlook for further research on the use of coastal dunes as a nature-based solution for coastal protection.

## I. Introduction

As climate change induces sea level rise and possibly heavier storms, coastal protection is in a transition phase from hard structural engineering towards soft measures that can adapt dynamically to a changing environment (Borsje et al., 2011; IPCC, 2014, 2018; Vousdoukas et al., 2018). Ecosystem-based approaches complementing engineering with functional parts of the natural system provide such an alternative to conventional coastal defense (‘hard engineering’). Indeed, estuarine and coastal soft sediment systems are dynamic by nature and their inherent ecological processes may be exploited to enhance resilience (Temmerman et al., 2013). Coastal foredunes represent the most important natural flood barrier for much of the European coastline and 30% of all shorelines worldwide (Martinez and Psuty, 2004; Reijers et al., 2019). In contrast to urban and other infrastructure, coastal dunes have the capacity to grow with rising sea level due to interactions between plant growth (Duarte et al., 2013) and aeolian sediment supply (de Vries et al., 2012; Strypsteen et al., 2019a). Therefore, they are currently considered as an important nature-based solution for coastal protection (Borsje et al., 2011; Duarte et al., 2013; Temmerman et al., 2013).

The use of foredunes as an engineering tool cannot be achieved without a deep understanding of the organizational properties of the natural dune system. Coastal dunes develop in first instance by sand accretion at the upper beach. In regions with predominant onshore winds, the magnitude of aeolian sand flux can primarily be described as a function of wind speed and grain size, but it also depends on soil moisture content, fetch length, beach and dune morphology (Delgado-Fernandez 2010, de Vries et al. 2012, Strypsteen et al. 2019). These aeolian fluxes impact the performance of a keystone species in foredunes from the European Atlantic coast : marram grass (*Calamagrostis arenaria (L.) Roth*, formerly *Ammophila arenaria;* Huiskes 1977). The species is the dominant species from white dunes, as protected within the directive 92/43 EEC [Shifting dunes along the shoreline with *Ammophila arenaria*, code 2120] (Perrino et al. 2014, European Commission 2007).

Biomorphogenesis refers to the process where biota like plants but also animals induce changes in the form of the environment they live in. Marram grass is such as an engineering species (Bakker, 1976; Puijenbroek et al., 2017) as its growth and performance depends on, and in turn influences, aeolian sand fluxes and hence, dune development (Hesp, 2002; Zarnetske et al., 2015; Strypsteen et al., 2019). Phenomena where the value of one state variable directly or indirectly affects the sign, direction and rate at which that variable changes, is defined as a feedback (Maxwell et al., 2017). These feedbacks can be positive (self-amplifying) or negative (self-dampening). As soon as sand dynamics cease, marram grass starts to lose its vigour and declines in abundance, making way for the development of grey dunes [called “Fixed coastal dunes with herbaceous vegetation (grey dunes)” code 2130* (European Commission 2007)] the next stage in the vegetation succession, dominated by drought tolerant mosses (Fig. 1). The degeneration of marram grass by sand stabilization was already noted by Marshall (1965), who called this phenomenon “The *Ammophila* problem” (not to be confused with the “Ammophila problem” referring to the invasion of the species outside its natural range as mentioned by e.g., Wiedemann & Pickart (1996).

**Figure 1.**
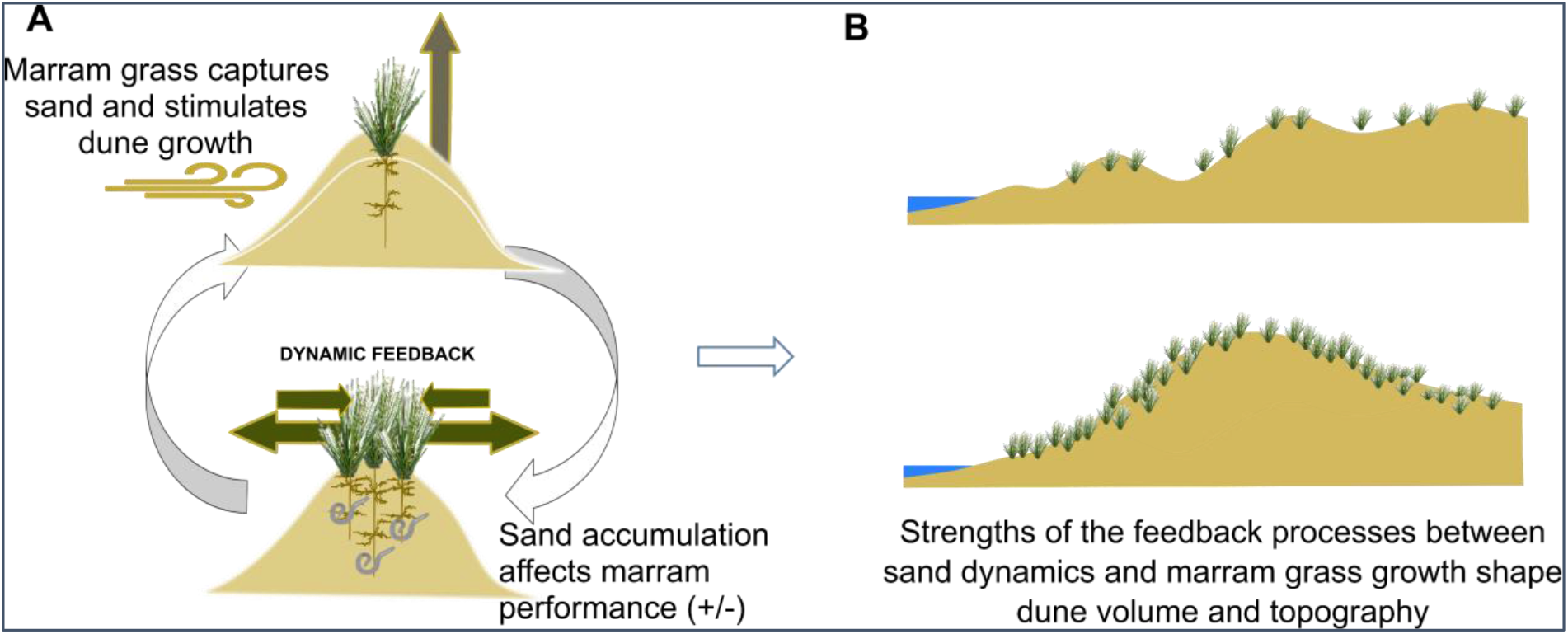
Conceptual figure on the role of marram grass as an engineer in foredune formation. A. Once established, the species’ sand capture ability shapes local sand accumulation, leading to an increase in dune volume. This sand accumulation will promote the species’ growth, unless burial is too severe. If sand accumulation ceases, either due to decreased input from the sea, or to sheltering effects from surrounding vegetation, the plant performance will decrease due to pathogen accumulation in the roots, after which marram grass will degenerate. B. These dynamic feedbacks depend on the species’ spatial configuration and external environmental conditions and will eventually shape the development of its volume and form, and, hence, its stability and resilience against storm surges under climate change.

The extraordinary sand fixing capacity of marram grass has been recognized in northwestern Europe for many centuries. *C. arenaria* was introduced for dune fixation in different parts of the world such as North America (Buell et al. 1995), Chile (Castro 1988), South Africa (Hertling & Lubke 1999), New Zealand (Hilton et al. 2004) and Australia (Webb et al. 2000). We here review the current state of the art with respect to the species’ biotic and abiotic drivers of performance (Part II). We subsequently review the quantitative evidence of feedbacks with sand dynamics and demonstrate by both a new model and remote sensing how marram grass spatial configuration affects dune development (Part III). We end this review by an outlook towards further research.

## II. Abiotic and biotic drivers of marram grass performance

### II.1 Abiotic drivers of marram grass performance

#### II.1.1. marram establishment

Marram grass establishes through seed germination or shooting of rhizome fragments detached from tussocks by coastal erosion. Konlecher & Hilton (2009) showed the potential for marine dispersal of such rhizome fragments over hundreds of kilometres, depending on regional sea currents. Seeds are mainly dispersed by wind, although the species only shows week morphological adaptations to wind dispersal (Huiskes 1979). Dispersal experiments by Pope (2006) and McLachlan (2014) suggest that a large majority of the seeds end up within a distance of less than 1 meter from the parent plant and wind dispersal abilities are probably restricted to several tens of meters. The potential for seed establishment of *Calamagrostis arenaria* is very high. First, this is due to a substantial seed production. Salisbury (1952) estimated over 20 000 caryopses are formed yearly per plant tussock. Second, marram grass has a long-lived seedbank. Viable seeds of up to 21 years old were recovered (Hilton et al. 2019). Third, the germination potential is high. Experiments under optimal laboratory conditions yielded germination percentages between 82 and 94% (Huiskes et al. 1979; Van Der Putten 1990, Bendimered et al. 2007, Lim 2011). In the field however, the establishment success of *C. arenaria* from seed is reputed to be very low on average (Huiskes 1977), although locally frequent germination (has been)/was observed in coastal dunes in The Netherlands (Van Der Putten 1990), New Zealand (Esler 1974) and North-America (Wiedemann 1987). Frequent establishment of marram is observed in embryonic foredunes and damp dune slacks (authors’ personal observations). Seed germination strongly decreases with sand burial (Van der Putten 1990, Lim 2011, McLachlan 2014). Seedling emergence decreases linearly with burial depth, with a 3 cm burial already resulting in a germination reduction of 60% and no more seedlings emerge when seeds are buried under 9cm of sand; Fig. 2). The results we obtained from a burial experiment (See Supplementary material 1) are very similar to the findings of Lim (2011).

**Figure 2.**
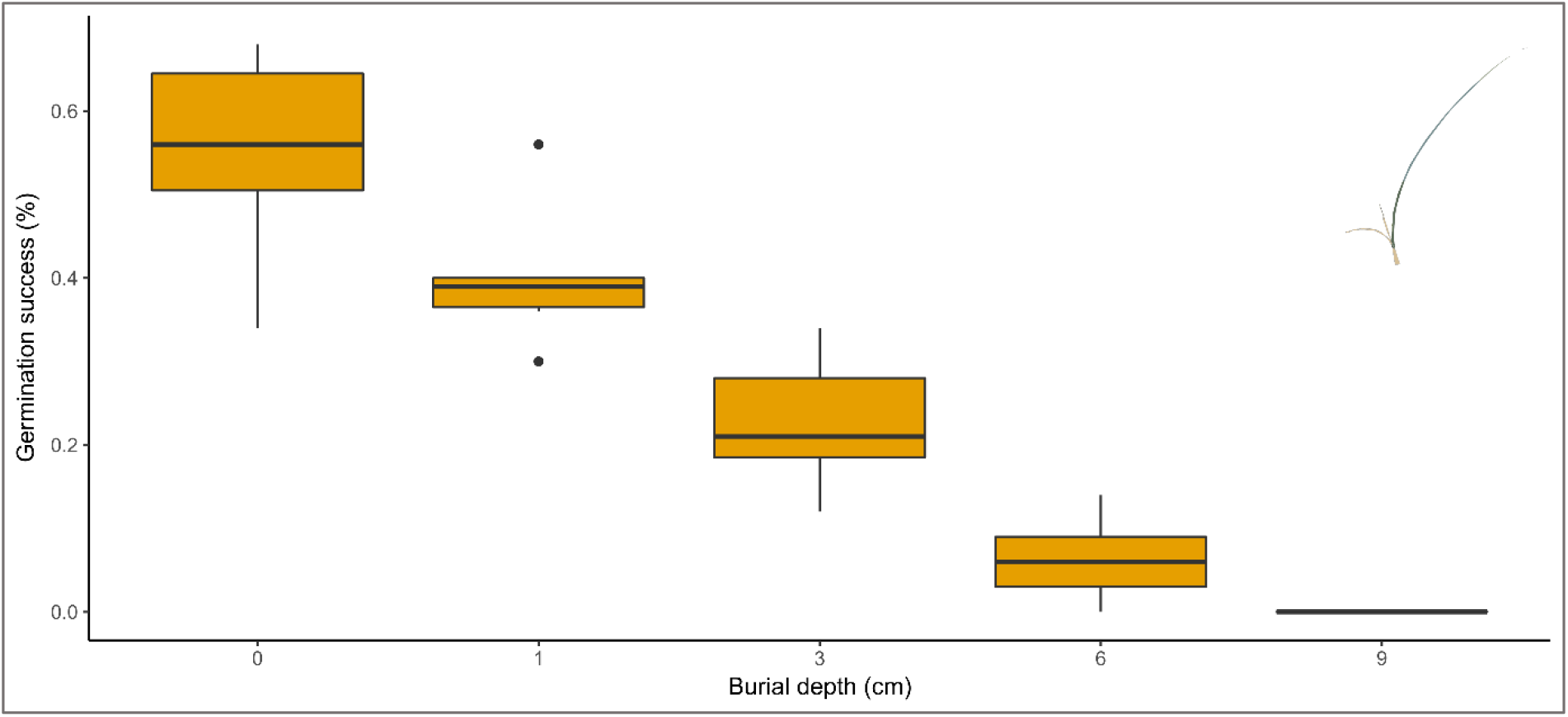
Seedling emergence in relation to sand burial (see supplement1)

Huiskes (1977) and Van Der Putten (1990) showed that the highest germination rates are obtained with a fluctuating (day/night) temperature regime and a day temperature exceeding about 20°C. These results are enhanced by stratification (cold pre-treatment). Optimal germination was obtained with a 20/30°C night/day temperature, with germination inhibited at lower temperatures of 10/20°C night/day. These germination requirements retrieved in the lab correspond well with observations of seeds germination in spring, when the temperature has risen sufficiently. Germination occurs only under moist conditions (Huiskes 1979) and is inhibited when salinity exceeds 9 g/l (Chergui et al. 2013).

#### II.1.2. Marram growth and survival

Once established, marram grass growth and survival depends largely on the exposure of its local environment for the physical forces of wind and water, that can directly dislodge plants or indirectly affect growth and survival by transporting sediment. Partial burial of seedlings resulted in a 50-60% increase of shoots length and root dry mass, but this vertical growth increases at the expense of lateral growth and overall shoot biomass (number of tillers, which was maximal at 0-40% burial of the shoot height) (Ievinsh and Andersone-Ozola, 2020). *C. arenaria* biomass increase showed a parabolic response to burial with optimal growing performance at burial rates of 31 cm of sand per growing season. (Nolet et al. 2018). The tolerance for burying was estimated to 78 to 96 cm burial/year. Reijers et al. (2021) found more mature tussocks (clonal fragments containing ± 8 shoots) to perform equally well under 0 or 2 cm burial every two weeks, but high mortality when burial reached 4 cm. Sediment burial also indirectly influence marram grass growth by protecting the plants against the detrimental effects of coastal flooding. Higher and larger embryo dunes are less susceptible to erosion during the winter storm season, which positively influences marram grass growth during summer (Van Puyenbroek et al. 2017).

Besides exposure to physical forces, soil nutrient levels can have a large influence on marram grass performance as well. In general, sandy coastal systems are nutrient-limited and *C. arenaria* can cope with these nutrient-poor conditions through symbiosis with arbuscular mycorrhizal fungi and by recycling its own plant material through slow decomposition (Kowalchuk et al. 2002, Reijers et al. 2020).

Despite its occurrence in nutrient-limited conditions, *C. arenaria* requires substantial levels of nitrogen, phosphorus and potassium for good growth (Willis 1965). A higher availability of N and P in lime- and iron-poor dunes, due to atmospheric deposition, has been proposed as a mechanism of the species’ local expansion in coastal dunes (Kooijman et al., 1998; Kooijman and Besse, 2002).

Increases in temperature, nutrients and precipitation stimulate vegetation growth and lead to a global greening of coastal dunes (Jackson et al., 2019). This global greening affects the natural sediment-sharing capacity of coastal dunes, by hampering sediment transport to the hinterland (Gao et al. 2020). Reduced sediment mobility and dune stabilization are thought to threaten several ecological functions, while it can increase the protective function of coastal dunes by lowering erosion susceptibility (Delgado-Fernandez et al. 2019, Gao et al. 2020, Pye & Blott 2020).

### II.2 Biotic constraints on marram grass performance

#### II.2.1. Negative plant-soil feedback

Marram grass was found to perform worse in its own rhizosphere soil than in either sand from the sea floor or in sterilized soil from its own rhizosphere (van der Putten et al. 1988, 1993), demonstrating that a biotic factor in the soil causes a decline in marram grass performance. The exact cause of this biotic control is to date unclear. The first studies attempting to pinpoint the soil organisms causing the decline of marram grass implicated root-feeding nematodes as well as pathogenic fungi (Van der Putten et al. 1990, De Rooij and van der Goes 1995, van der Putten and van der Stoel 1998, van der Stoel and van der Putten 2002). However, the exact species causing a performance reduction could not be identified across these studies. Competitive and facilitating interactions among these co-infecting belowground parasites (Brinkman 2005&a,b,c) but also more complex trophic interactions, including those with microbes within the rhizophere (Piśkiewicz et al. 2008, 2009; Costa et al. 2012) were found to be mediators of marram performance under experimental conditions. Furthermore, it has been shown that the negative effect of certain nematode species can be mitigated by the positive effect of mycorrhizal and endophytic fungi (Little and Maun 1996, de la Peña et al. 2006, Hol et al. 2007). Overall, the net effect on marram grass performance of all naturally occurring members of the soil community is generally negative. Although the exact mechanism is difficult to identify, evidence for the “escape hypothesis” remains strong, i.e., marram grass needs regular burial by wind-blown sand free of soil organisms so that it can grow new roots into an –at least temporarily– enemy-free space. Plant-soil feedbacks caused by other plant species also play a role. Conditioning of soils by *Carpobrotus edulis* (L.) N.E.Br., a species originating from South Africa and one of the most invasive plant species in the Mediterranean, suppresses marram grass biomass and in some cases survival rate (de la Peña et al. 2010). The increase of *Carpobrotus* in the dunes of central Italy (Sperandii et al. 2018) has therefore been linked to large-scale decreases in marram grass. Marram grass also shows a reduced germination on soil invaded by *Acacia longifolia* (Andrews) Willd., yet it performed better on invaded than native soil after 12 weeks of growth (Morais et al. 2019).

#### II.2.2. Aboveground biotic interactions

The aboveground organisms associated with marram grass are in general well known (i.e., Huiskes 1979, Heie 1982, 1986, Holman 2009, Vandegehuchte et al. 2010a), but very little is known about their effects on plant performance. The associated herbivore species have the potential to induce serious reductions in aboveground performance in a controlled environment (Balachowsky and Mesnil 1935, Nye 1958, Heie 1986, Vandegehuchte et al. 2010a), but so far no experiments were conducted in nature. Marram grass does not seem to be controlled to any significant extent by mammalian grazers either (Badhresa 1977, Huiskes 1979), except for some feeding on young shoots (Rowan 1913). Seed predation has been observed (Huiskes 1979) but its magnitude and/or impact on marram grass demography is unknown.

#### II.2.3. Control of above- and belowground communities by marram grass intraspecific variation

Intraspecific variation among marram grass populations can have strong effects on the abundance and community composition of both above- and belowground invertebrate species (Vandegehuchte et al. 2011). This variation is linked to genetic variation in plant growth, which likely explains higher abundances of aboveground invertebrates on local than on non-local marram grass populations. Contrasting effects were found for root herbivores as their abundance and species richness negatively covaried with the aboveground ones (Vandegehuchte et al. 2011, 2012). Additionally, it has to be noted that a full soil biota community can have stronger effects on marram grass performance than local abiotic soil properties (Vandegehuchte et al. 2010c), although performance can differ significantly among soils differing substantially in abiotic properties. The relationships between marram grass and its aboveground invertebrates can therefore not be understood independently of its belowground invertebrates and the abiotic conditions of the soil.

#### II.2.4. Learning from elsewhere: marram grass as invasive species

Explanations for success of marram grass in its novel range have been sought in the popular “enemy release hypothesis” (Keane and Crawley, 2002), mainly focusing on belowground enemies. Growth of marram grass was significantly less reduced on soils from South African sites than on soils from the Netherlands, indicating a weakened negative plant-soil feedback and thus potential role for enemy release in South African soils (Knevel et al. 2004). However, this contrasts with findings from coastal dunes of California, where soil sterilization experiments have shown that the performance of marram grass is reduced to similar extents as in Europe when grown on non-sterilized soil (Beckstead and Parker 2003), suggesting there is no enemy release. Furthermore, soil biota from three native South African plant species did not suppress marram grass growth, but biota from soils beneath the tropical cosmopolitan dropseed *Sporobolus virginicus* (L.) Kunth did, suggesting that this plant species may confer biotic resistance against invasion by marram grass (Knevel et al. 2004). A large sampling campaign of soil and roots from Tasmania, New Zealand, South Africa and the west coast of the USA revealed that marram grass did not have fewer root-feeding nematode taxa in these regions than in its native range. However, native plants in the novel range had more specialist root-feeding nematode taxa than marram grass, while specialists such as cyst and root-knot nematodes, which are common in the native range of marram grass, were not found in the southern hemisphere (van der Putten et al. 2005). Invasiveness of marram grass thus seems correlated with an escape from specialized root-feeding nematodes.

### II.3. Dynamic feedbacks between aeolian fluxes and vegetation development

The capture rates of sand by vegetation and its effect on dune topography have been intensely studied (e.g., Hesse & Simpson 2006). There is also abundant literature on how obstacles that represent vegetation obstruct or facilitate sand fluxes, with strong analogies to research on fluid dynamics. Typically, multiple configuration of height and density of the obstacles are used (e.g., reed stems (Arens et al., 2001); see (Mayaud and Webb, 2017) for a comprehensive review on aeolian sand transport in drylands). These studies quantify how much of the total force of the wind by drag is reduced by the vegetation, also referred to as shear stress partitioning and expressed as drag coefficients (Raupach, 1992). All studies show this drag coefficient to be positively related to the roughness induced by the density and impermeability of the set of obstacles, and their height (Hesp et al., 2019). Since these experiments use marram-grass surrogates like artificial cylinders, stem bundles or even dead plant material, they do not represent the realised morphology of dune vegetation, which precludes further progress in understanding the feedbacks between sediment capture and plant growth. Clusters of tillers enhance sand deposition by lowering wind speed and associated shear stress within the vegetation canopy (Charbonneau & Casper 2018). Larger tussocks are able to capture more sand, thereby imposing a positive feedback on their own development and vigour. The plants react to burial by rapid production of elongated stem internodes, but the exact extent of this growth response is unknown except for young plants under lab conditions (Levinsh and Andersone-Ozola 2020). As sand burial induces the production of high-density vertical tillers and horizontally expanding rhizomes (Reijers et al., 2021), marram grass steers dune morphology (van der Putten et al. 2005, Hart et al. 2012, Darke et al. 2016). *C. arenaria* is, because of this growth strategy, associated with the development of higher and steeper dunes compared to those formed by its North-American sister species *C. breviligulata*, making dunes build by the former potentially more resistant to erosion (Zarnetske et al., 2012; Seabloom et al., 2013; Charbonneau et al., 2016).

Both vertical and horizontal growth responses influence the size and shape of *C. arenaria* tussocks, but also directly determine remaining sand drift at the rear side of these vegetated patches (Reijers et al., 2021). With increasing densities and cover, *C. arenaria* subsequently stabilizes the mobile sand (Huiskens 1979). At least in European coastal dunes, the ceasing sand fluxes mediated by the species’ increasing densities, and the resulting increases in dune height and slope, then induce on longer time frames a negative feedback on the species’ vigour in the long run, causing the species to slowly die off (e.g., Van der Putten 1994, De Rooij and Van der Goes 1995, van der Putten and van der Stoel 1998, van der Stoel and van der Putten 2002) as the resource (fresh sand) becomes limiting. The spatial configuration and morphology of the vegetation is therefore dynamically coupled to shear stress. Sand capture directly alters potential density, growth and lateral expansion of the vegetation, which feedbacks to patterns in flow parameters (velocity, turbulence, intermittency) because of sheltering effects by vegetation and dune topography. The qualitative importance of these feedbacks for the large-scale geomorphology of coastal dunes is well-appreciated (Durán and Herrmann, 2006; Hesse and Simpson, 2006; Durán et al., 2009; Durán and Moore, 2013), but very few data on the feedbacks between sand fluxes and vigour of the foredune vegetation are available. So far, different vegetation states rather than vegetation dynamics have been linked to dune height potential and subsequent risks of overtopping events and flooding (Seabloom et al., 2013).

## III Towards an integrated model of the plant-sand feedbacks

### III.1 The integration of vegetation-dune feedbacks in existing process-based models

The importance of the vegetation-dune feedbacks is still not well understood, let alone quantified and incorporated into predictive models for coastal dune dynamics (Puijenbroek et al., 2017). Current state-of-the-art 3D models for coastal dune development (e.g., DUBEVEG (Keijsers et al., 2015), CDM (Durán and Moore, 2013), AeoLIS (Hoonhout and de Vries, 2016)) are able to simulate topographic development of coastal dunes and sediment transport at spatial scales relevant for coastal managers as a function of sediment supply, probabilities of vegetation development, descriptions of flow field, and dune erosion by waves. These (coupled) 3D coastal dune models are a product of the basic physical principles and sediment transport models, and they are essential for the prediction of dune development. Furthermore, they need validation from field experiments containing high-quality datasets relevant for dune development. With exception of the DUBEVEG model (De Groot et al., 2011), which has coarse vegetation dynamics incorporated, recent coastal functioning models have ignored ecological interactions across scales (e.g., Duran and Moore 2013, Van Westen et al. 2019). As important engineer species from coastal dunes differ in physical features and life history, they differently affect dune development, with for instance sand couch grass (*Elymus farctus* (Viv.) Runemark ex Melderis) giving rise to ‘lower broader dunes’, and marram grass enabling dunes to develop into a ‘higher, hummocky peaked topography’ (Hesp, 2002; Puijenbroek et al., 2017; Reijers et al., 2019; Schwarz et al., 2019).

All existing dune erosion models treat the processes at the dune face in a simplified way. No process-based description is implemented to describe the formation of vertical cliffs at the dune foot undermining the dune slope with subsequent geotechnical failures of the dune slope that results in slumps of sand on the beach that can be taken away by the waves. For this marshes, Bendoni et al. (2019) implemented a hydro-morphodynamic interaction model in XBeach (Roelvink et al., 2009) to evaluate erosion of marsh boundaries due to wave impact. Although this study is limited to the cohesive sediments’ environment, soil reinforcement due to roots has been modelled, which might be extended to other environments in the future. Physical scale model experiments, with dunes, vegetation and disturbances scaled towards lab conditions, have demonstrated that roots, which geotechnically strengthen a sand volume, significantly reduce the dune erosion compared to bare sand (Feagin et al. 2019, Bryant et al. 2019). Presently, only indirect implementations are possible namely by tuning calibration parameters influencing the morphodynamics.

### III.2. Insights from a new simulation model

#### III.2.1. The geography of marram spatial configuration

From II.3, it is clear that feedbacks between the environment and the spatial distribution of marram grass impact dune development. We mapped marram cover and spatial contingency in 20×20 m^2^ grid cells along the coastlines of northern France, Belgium, Netherlands and South-England (see supplement S2), to identify realistic ranges in nature. Marram grass is –as predicted from the species’ biology-predominantly showing a clustered distribution with JC (join-count; an established method that quantitatively determines the degree of clustering or dispersion, see S2) values between 20 and 80, so ranging from random (values close to zero) to highly clustered patterns (Figure 3). A mean clustering pattern with JC values around 50 is stable over the four studies regions. No underdispersed (so regular) patterns were observed. Interestingly, marram grass spatial cover at these spatial scales is strongly country-specific with UK and France being represented by well-vegetated dunes. Dunes in Belgium and the Netherlands appear to be in more dynamics states with quite a substantial presence of areas with a low cover (see IV).

**Figure 3.**
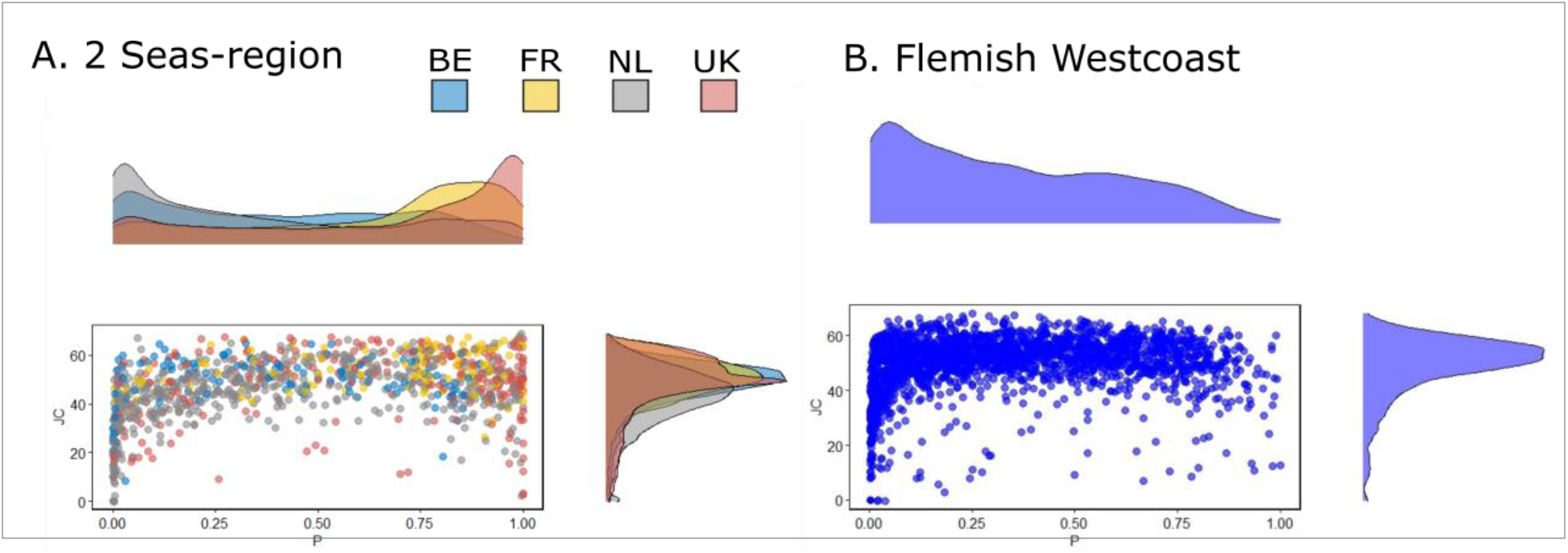
Density distribution plots of observed cover (P) and spatial correlation (JC) of marram grass in 20×20 m^2^ grid cells in dunes from the Isles of Scilly to Norfolk in England (UK) and from Somme (Fr) to Texel (NL) on the continental side along the North Sea and the Flemish West coast (B) (right panel). Note that for visualisation, a subsampling of 1000 points (2%) was performed for (A).

#### III.2.2. Simulating dune growth in relation to marram grass spatial configuration

We developed a grid-based dune simulation model that computes aeolian transport processes and changes in vegetation growth and dune morphology based on their dynamic feedbacks and marram spatial organisation. The landscape is grid-based with cells having dimensions of 0.20×0.20 m^2^. The 100×100 cell matrix therefore corresponds with a dune area of 20×20m^2^. We refer to supplementary material S2 for a detailed process overview, references to the code. We simulated changes in aeolian processes and wind dynamics at a day-resolution. We used averages over 7 years (2010-2017) received from the Royal Meteorological Institute at Koksijde, and scaled them to the four main different wind directions as used in the model by (Nolet et al. 2018b), each corresponding to a side of the landscape. Wind speed and direction is drawn daily from a normal distribution, based on monthly average wind speed and its standard deviation. *Sand input*, the material blown into the system from the beach (N-direction here), is expressed as a relative percentage of the maximum sand saturation flux, i.e. the maximal amount of sand that can be carried by the wind. Lateral winds (E and W-directions in the grid) have an influx which corresponds with the most recent outflux of a lateral wind (thus, we represent the landscape as tube to avoid edge artefacts). This amount is constantly updated during a simulation.

*Sand deposition* is directly dependent on shear velocity which is a function of wind velocity Hoonhout and de Vries 2016; Durán et al., 2010), vegetation density (Durán et al. 2010) and its roughness (Durán and Moore 2013). Increases in shear stress due to funnel effects are included, as are gravity and shelter effects. Maximum angles of repose are set to 34° (Durán et al., 2010)when vegetation is absent (Durán et al., 2010). These angles increase with vegetation density. As such, avalanches are less prevalent when plant density is high. Moreover, erosion is inhibited in locations sheltered by lee slopes at an angle of maximal 14° (Kroy et al., 2002). *Marram grass* dynamics are seasonal (only growth in spring and summer), with local growth modelled as outlined in this review. Vertical as well as lateral growth during the growing season is modelled as a logistic function up to maximal heights, and directly depending on sand deposition (Nolet et al., 2018), leading to positive growth under intermediate sand accretion and complete burial leading to marram grass die-off. Lateral growth follows Lévy-patterns as determined by (Reijers et al., 2019), and are here modelled by simplified neighbour expansion processes. No growth occurs during autumn and winter but sand accumulation continuous. The net height after winter burial determines the starting conditions for vertical growth in the next season. No germination events were modelled as these are to date not (or only rarely) witnessed in foredunes the last decade.

To validate the model predictions, we compared outcomes from the model with those from a statistical model linking changes in topography over five years as derived from LIDAR in relation to the initial marram spatial configuration as determined from aerial photographs in 2015 in Belgium (Fig. 4; detailed methods in supplementary material S3 and S4). Our model simulations (Fig 5, upper panels) were run for initial marram cover in the range 0.1-0.9 (as without marram cover, only erosion of the initialised sand volume is occurring without establishment) and join counts between 20 and 70).

**Figure 4.**
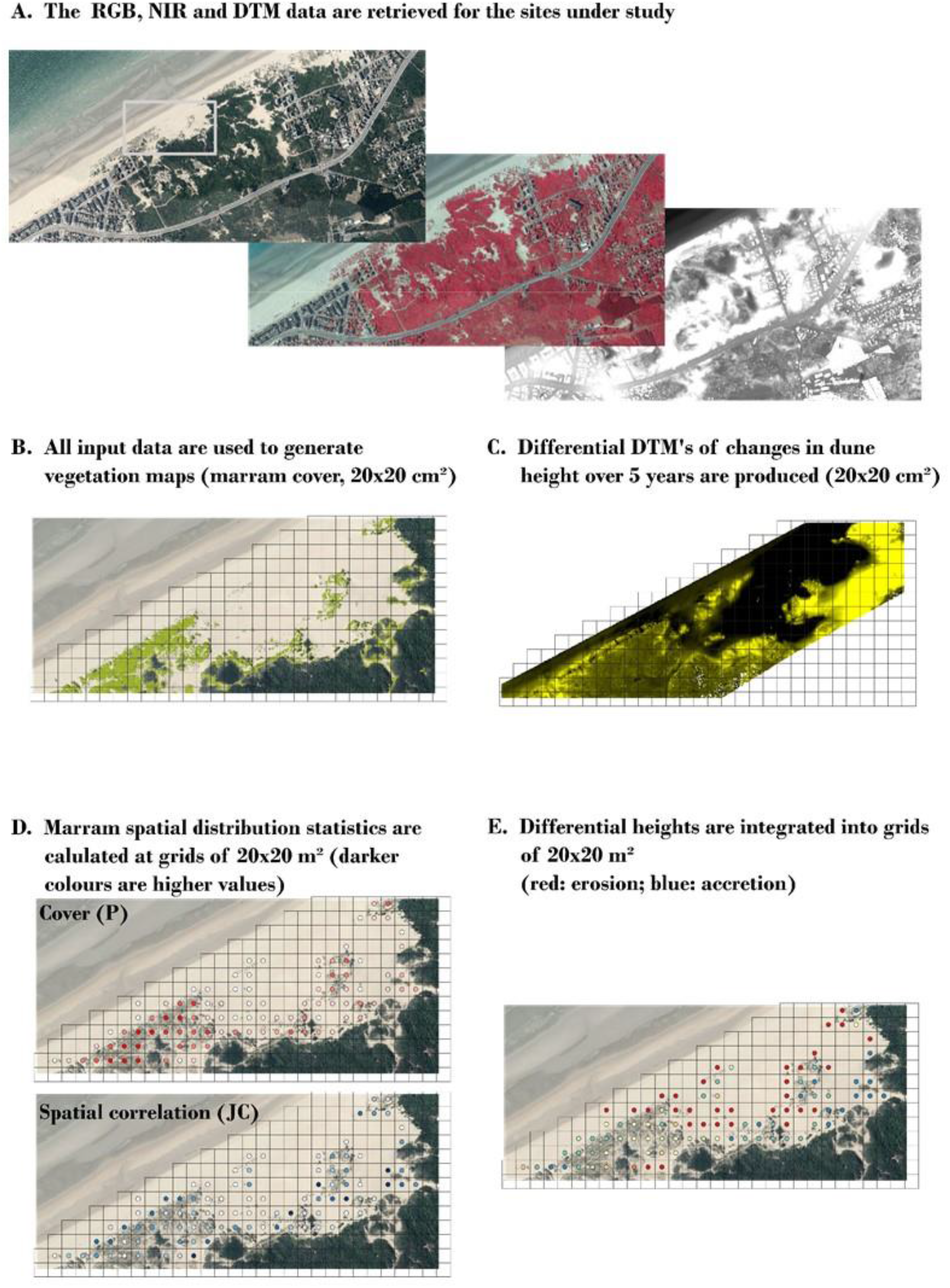
Workflow for connecting observed differences in dune height over a period of 5 years to marram spatial configuration in coastal foredunes (case presented: Schipgatduinen, Koksijde, Belgium)

**Figure 5.**
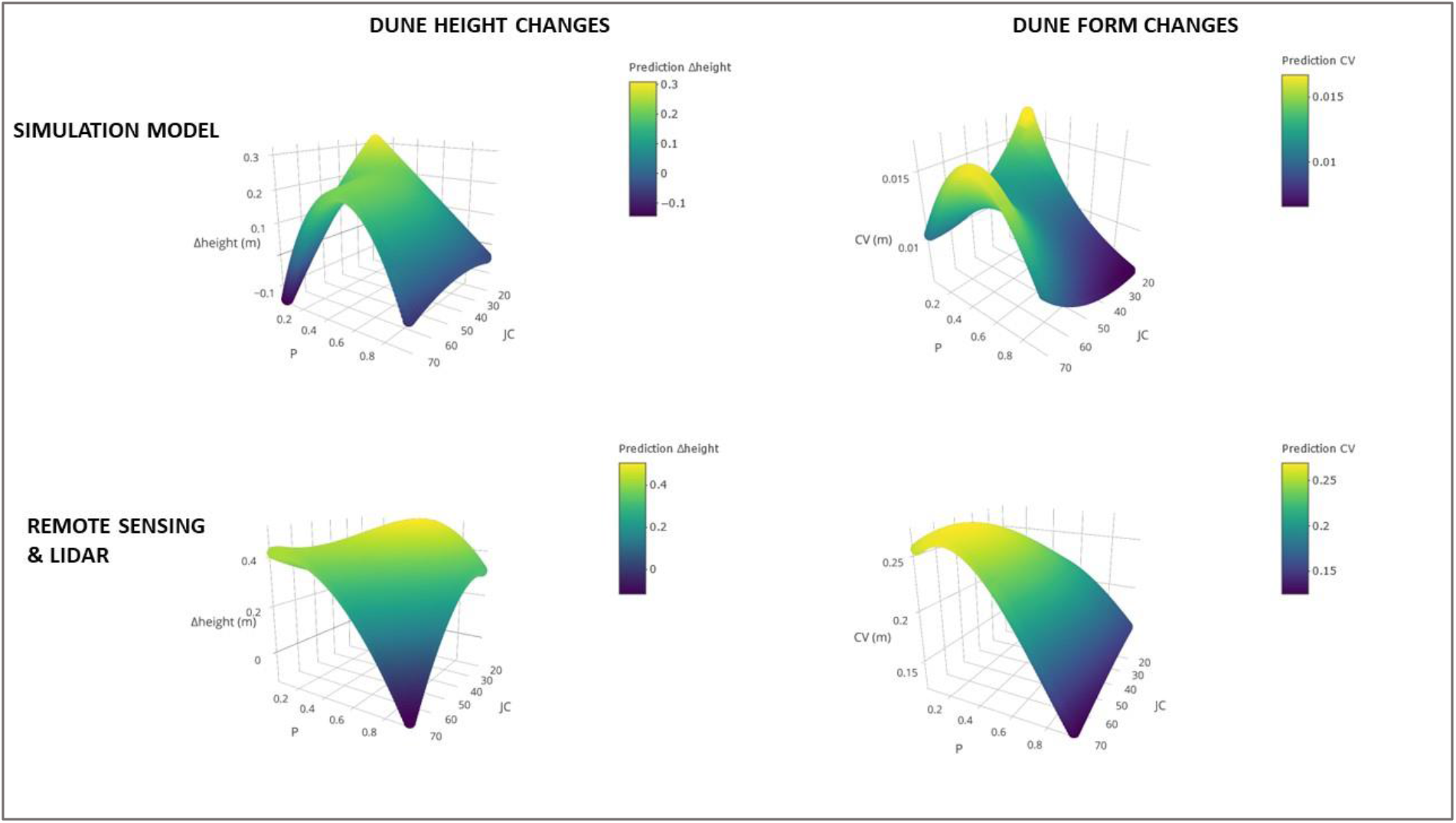
Output of the simulations computing changes in dune height and topography (CV of height changes over all grid cells) over a period of 5 years (upper panel), and similar metrics as observed from LiDAR data from coastal dunes along the Belgian west coast (lower panels).

The modelled height changes agree in general terms with those observed. The observed larger effects in the field therefore suggest slightly larger sand input, either due to sand availability or changes in wind strengths, from the beach as sand input initiated in the model based on rough estimates from the Belgian coast (unpub. data Rauwoens & Strypsteen). Alternatively, the small scale of the mechanistic model may underestimate wind saturation and therefore sand displacements (see IV. outlook). Dune height increases at intermediate cover of marram grass, so *P~0.5*. However, the simulation model predicts increases to be maximal under low cover and more random (i.e., less clustered) distributions of the marram grass tussocks (low *P* and low *JC*). According to the simulation model, local changes in dune topography, estimated as the coefficient of variation (CV) of grid cell-level differences in height, show most changes occurring in dunes with clustered marram patches or patches with a low amount of vegetation, but more random patterns. Predicted changes from LIDAR follow the same pattern as the ones generated by the simulation model. Only under low cover and intermediate clumping, a more homogenous increase of the dune is predicted by the computer model.

Analysis of the LIDAR data also showed decreasing dynamics with increasing distance from the sea (see supplement S4). The obtained effect sizes (supplement S4) and accompanying visualisations of the modelled effects (figure 5) indicate that the observed changes in integrated dune height and form differ from those of the simulation in the sign and strength of the cover x spatial clustering (P:JC and the interaction between P^2^ and JC). The most prominent difference lies in the predicted erosion dynamics under low marram cover and a strong clustering. This suggests that the sand accretion capacities of marram grass under these conditions needs to be re-evaluated.

## IV. Discussion and outlook

A mechanistic understanding of the vegetation-sedimentation feedbacks that steer the natural development of coastal dunes is essential for conserving and restoring the function of coastal dunes as natural flood barriers. Climate change, and its impact on feedbacks between marram grass and sand fluxes, is anticipated to strongly alter dune formation and dune resilience (Pakeman et al., 2015; van Puijenbroek et al., 2017). We here reviewed the state of knowledge on the ecology of marram grass in relation to dune formation, flow attenuation, sediment deposition and plant growth. Our model and LIDAR analysis showed that the joint increase of volume and variability under low cover and less clustered spatial configurations have the highest impact on local sand accretion and dune morphology. Such conditions steer impose a positive feedback on vertical growth. Strong erosion dynamics are conversely anticipated to preclude establishment at further distances from existing tussocks. Scale-dependent feedbacks lead to patterns of self-organisation (Rietkerk & van de Koppel 2008) and need to be quantified and further integrated into mechanistic models to forecast coastal dune formation in relation to climatic conditions. Earlier research showed considerable variation in marram growth (Vandegehuchte et al. 2010a) and expansion strategies (Reijers et al. 2021), and changes here-in can be expected with respect to future climatic conditions. The relevance of this intraspecific variation remains to be understood, also from the perspective of planting actions to actively build resilient dunes in the light of climate change.

A resilient coastal dune system is anticipated to be one where vegetation and bare sand coexist in a stable equilibrium, hence a state to which the system should bounce back after any disturbance, e.g., by erosion. The permanent loss of sand dynamics by changing sand input, fragmentation or anthropogenic dune stabilization are expected to lead to catastrophic shifts causing dunes to become hyperstatically fixed by plantation and succession (Jackson et al. 2019, Gao et al. 2020). On the other hand, at too low initial densities, the vegetation may be disrupted by strong sand drifts, also following intense trampling by people, leading to a hyperdynamic and unvegetated state. A resilient dune should balance between both extremes (Borsje et al. 2011) and this resilience will thus largely be determined by the current vegetation density and configuration, local conditions of sand supply, connectivity with the beach, and erosion. The state of the marram dune can be expected to impact further inland sand drift. Narrow stretches thereby have the potential to determine dune stability at larger spatial scales by affecting the total dune system volume, and the further vegetation succession dynamics (e.g., Ollf et al. 1993, Fenu et al. 2013). These are less relevant from a coastal protection perspective but of major importance for biodiversity conservation (European Commission on Habitat of Directive 92/43 EEC)

Coastal dunes along the coast of the North-Sea and Channel show a remarkable convergence in the spatial clustering across the four studied countries, and this clustering seems to be preserved across the range of vegetation cover. This finding suggests an optimal clustering in European dunes, which is anticipated to result from the species’ self-organizing capacity. At intermediate cover, this clustering leads to largest changes in dune growth. We anticipate that the availability of sufficient aeolian dynamics at small scales drives this overall increase in dune volume. This review also shows these conditions to facilitate marram grass performance because of the steady supply of fresh sand. Although more research is needed, this finding suggests that such a spatial configuration can optimize both marram grass performance and dune resilience by maximizing growth. Deviations from this state, especially in terms of cover –note that the clustering metric becomes less relevant with increasing cover– are then likely disturbed states resulting from either ceasing sand dynamics or vegetation development. As we only documented patterns in marram grass spatial configuration, we lack insights into the underlying causes. Are they due to sediment transport potential, or correlated to region-specific variables such as tidal amplitude, wave height, or beach width? Alternatively, it is not unlikely that variation in both dune management –especially planting campaigns– and the differences in recreational pressure are at the basis for this variation. Since we showed marram grass’ spatial configuration to affect both dune growth and topography and therefore sand fluxes further inland, this variation is anticipated to have strong implications for coastal protection. Pending on the state, recreational pressures may constrain dune stabilisation and keep the system in a dynamic, and presumed optimal state with respect to resilience, or facilitate erosion and the transition to hypermobile states. Clearly, the negative and positive contributions of such recreational pressures need to be assessed case by case, and in direct connection to the local environmental (boundary) conditions (Nunes et al. 2020).

Incorporating the available information allowed us to mechanistically build models that support generic predictions of dune volume and topography change at short spatial and temporal scales. The model does, however, still contain gaps in terms of parameterisation and validation (both observational and experimental), especially with regard to very dynamic conditions (low, clustered cover by marram grass). While any prediction in this specific parameter range can be questioned for its relevance (“How natural are these configurations, if we do not observe them?”), we argue that this is of the utmost relevance in the light of dune-building campaigns where marram grass needs to be planted, for instance in front of existing dikes. The presented simulation model also operates at relatively small scales relevant for vegetation dynamics, but potentially underestimating realised wind saturation in barely vegetated dunes. Deviations between the observed and predicted changes in dune volume likely result from such scaling issues. Upscaling of dune-vegetation dynamics can be achieved by linking sand-output conditions from the most seaward-oriented dunes as input conditions for those more inland. Under such conditions, sand-vegetation dynamics need to be extended towards other species occurring along the expected succession gradient. It remains to be studied whether simplifications using vegetation height and biomass (Duran *et al*. 2009), as the mediator of such interactions suffice. A major driver of dune development is the amount of sand input from the beach. While difficult to measure, the joint analysis of vegetation and dune development may be used as a reliable predictor of such sea-land sediment fluxes by means of inverse modelling.

More research is needed on potential regional changes in the interactions among vegetation development, dune growth and sand fluxes, for example caused by differences in climatic conditions and marram grass genetic variation. Dune volumes are prime determinants of their functioning in coastal protection, with large bodies of sand providing a larger safe zone against inundation risks from storm surges. Vegetation dynamics are expected to have a strong impact on the healing capacity of foredunes, i.e., how fast they recover to earlier states after erosion by such storm events, or in the longer term resulting from sea level rise. Current models enable us to predict changes in dune volume and form, but it remains unknown how erosion is affected by the vegetated state of the dune. For example, to which degree erosion resistance is determined by the marram grass root network still needs to be elucidated. Ultimately, the addition of such information on the mechanistic underpinnings of the patterns generated by our models would reduce the uncertainty of their predictions to the benefit of all stakeholders involved.

## Contribution to the Field Statement

Coastal dunes provide safety against future climate change induced sea-level rises and storm surges. Contrary to hard protections, they have the advantage to grow with the sea-level. Coastal dunes thus provide important safety services, especially in highly urbanised lowland areas. Because these coastal landscapes are formed by the interplay between the dune building marram grass and sand dynamics, a quantitative study of these interactions is necessary to understand and forecast how climate change will impact dune development and its resilience. We here provide a synthesis on the available insights, and complement these insights with new experiments and remote sensing data from coastal dunes along the North Sea. This information is integrated into a mechanistic model to demonstrate the importance of marram grass ecology, and its emerging spatial configuration for dune development.

We build on these insights to identify important caveats and future research directions to effectively manage and construct dunes as a solid nature-based solution from the fundamental ecological and geomorphological building blocks.

