## supplementary material for "Biomorphogenic feedbacks and the spatial organisation of a dominant grass steer dune development"

### *Supplementary material 1: methods for burial experiment*

An experiment was carried out to investigate the ability of *Calamagrostis arenaria* seed to germinate and emerge from sand burial. *S*eeds were harvested in august 2013 and sown in January 2014. Seeds were stored dry at about 20°C except for a 3 week stratification treatment at 5°C. They were subjected to four (1, 3, 6 and 9 cm) burial treatments. A control where seeds were sown superficially was added. Each treatment was replicated 5 times and for each replicate 50 seeds were used. The seeds were sown evenly spaced in pvc tubes with 10 cm diameter filled with 10 cm of a sand and garden soil mixture. The latter was used in order to prevent rapid dehydration of the substrate. The bottom of the tubes was sealed with a geotextile enabling drainage of excess water. Finally, layers of mineral dune sand with thickness according to the treatments were added on top of the seeds. Treatments were watered till saturation twice a week, simultaneously with seedling counting.

### *Supplementary material S2. Quantification of the Calamagrostis arenaria spatial configuration*

#### S2.1 Acquisition of the remote sensing data and indication of dune areas

The study area covers the coastal dunes of the United kingdom, France, Belgium and the Netherlands (figure S2.1). A different source of remote sensing data had to be used for each country. However, we searched for aerial images of similar quality between countries and selected images with minimal cloud cover and shadow interference. The aerial images differ in origin, in resolution and in the year in which they were taken (table S3.1). Aerial photographs were combined to obtain the RGBI band: blue (452–512 nm), green (533–590 nm), red (636–673 nm) and NIR (near infrared, 851– 879 nm). Digital Terrain Models (DTM) and Digital Surface Models (DSM), derived from LiDAR data, were obtained where available (Belgium and the Netherlands, table S3.1). For each country, the dune areas were mapped manually by visually interpreting the aerial photographs. The mapping was done with QGIS (QGIS Development Team 2020). Dune areas were considered up to 200m land inwards for France, the Netherlands and the UK. Because the vegetation maps for Belgium are used in other projects as well, the dune areas were mapped further landward (up to 3.8 km).

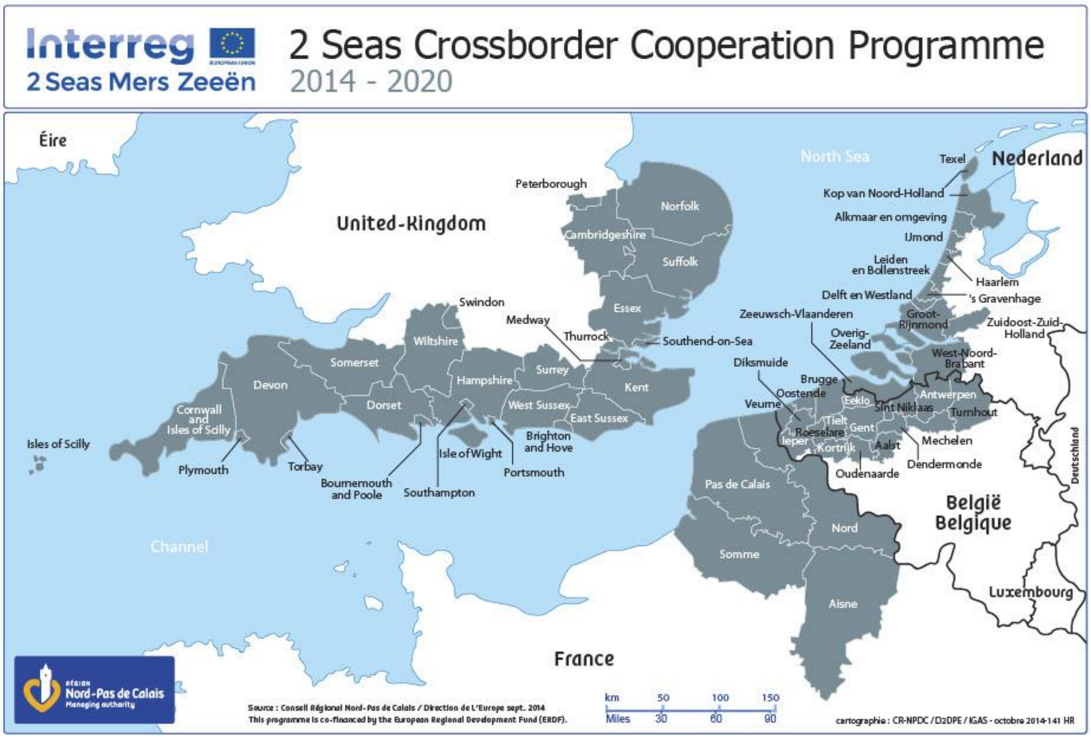

*Figure S2.1: The Interreg 2 Seas region. Aerial photographs were gathered for all indicated coastlines.*

**S2.2 Classification**

Support vector machines (SVMs) are a supervised non-parametric non-statistical learning technique (Lucas 2020) for which no assumptions are made on the underlying data distribution. The SVM is trained by making use of labelled data for which the algorithm aims to separate observations into a number of predefined classes. This trained algorithm is then used to classify all the data into the classes matching the training set (Mountrakis *et al*. 2011). We decided to use a SVM because of its efficiency at generalizing despite using a small training set (Mountrakis *et al*. 2011).

Vegetation maps are derived from the remote sensing data by classifying every pixel within the indicated dune areas into one of the expected vegetation types using a SVM. The expected vegetation types are: trees, shrubs, heath, grassland, marram grass, grey dune & sand. The used classes differed between the countries since some vegetation types did not occur in all countries (table S2.2). As a result one model was constructed per country. For each vegetation type, polygons containing pixels of the same type were drawn manually on different images. Half of the pixels was used as reference data for training while the other half was used for validation of the classification The same number of pixels was used per vegetation class for training of the SVM within one country, however, in order to have a sufficiently high accuracy of the resulting classification, the number of pixels did differ between countries. The SVM's use the 4 bands described earlier (RGBI) together with the Normalized Difference Vegetation Index (NDVI), which can be calculated from band 3 & band 4 as NDVI = (NIR-RED) / (NIR+RED) (Tucker 1979; Pettorelli 2013). The height of the vegetation, calculated from the LiDAR data by subtracting the DTM from the DSM, was also incorporated to further optimize the classification.

Apart from validation by the algorithm itself, the classified vegetation maps were also validated through ground truth data of specific areas that were not integrated into the training dataset. If the ground data showed that the model was not effective enough in classifying some vegetation types, extra training (via inclusion of extra pixels) of the algorithm was done to improve the outcome.

In the end, all four classifications had an overall accuracy higher than 90%. However, the models for the UK and France had a higher accuracy and kappa than those for Belgium and the Netherlands (table S2.3). The confusion matrix performance on the training dataset, together with the predictor accuracy and the user accuracy (two measures for the ratio of correctly classified pixels to the total number of truth pixels; Foody 2002, Thoonen *et al.* 2008) can be found in table S2.4.

All SVM's were constructed using the package "e1071" (version 1.7-2, Meyer *et al.* 2019) in R (version 3.6.1, R Core Team, 2019), all with a Radial kernel (gamma = 1/6, cost = 1). Since the LiDAR data had a lower resolution than the colour images (1m and 0.25m, respectively), we rescaled the data before including them in the SVM models. This rescaling was done using resampling with nearest neighbour in the package "raster" (version 3.4-5, Hijmans 2020).

*Table S2.1: Origin of the remote sensing data per country.*

| **Ortho-photos** | **The United Kingdom** | **France** | **Belgium** | **The Netherlands** |
|---|---|---|---|---|
| Source | Airbus | Airbus | Flemish government | Dutch government |
| Origin | Pleiades satellite | Pleiades satellite | Planes | N° of different satellites |
| Year | 2016-2019 | 2018 | 2015 | 2018 |
| Link + description | Website of Airbus https://www.airbus.com/ | Website of Airbus https://www.airbus.com/ | Orthophoto mosaic mid-scale summer shots 2015 Flanders https://download.vlaanderen.be | Orthophoto composed with images from different satellites – 2018 https://data.nlextract.nl/beeldmateriaal/2018/ |
| Resolution (m) | 0.5 | 0.5 | 0.4 | 0.25 |
| Raster dimensions (m) | 25x25 | 25x25 | 20x20 | 25x25 |
| N° of pixels per raster cell | 2 500 | 2 500 | 2 500 | 10 000 |
| CRS* | British National Grid - EPSG:27700 https://epsg.io/27700 | WGS 84 / UTM zone 31N - EPSG:32631 https://epsg.io/32631 | Belge Lambert 72 - EPSG:31370 https://epsg.io/31370 | RD New - EPSG:28992 https://epsg.io/28992 |

| **LIDAR** | | | **Belgium** | **The Netherlands** |
|---|---|---|---|---|
| Source | | | Flemish government | Rijkswaterstaat |
| Year | | | 2015 | 2018 |
| Link | | DSM:<br>DTM: | https://download.vlaanderen.be/Producten/Detail?id=937&title=Digitaal_Hoogtemodel_Vlaanderen_II_DSM_raster_1_m<br>https://download.vlaanderen.be/Producten/Detail?id=939&title=Digitaal_Hoogtemodel_Vlaanderen_II_DTM_raster_1_m | https://www.pdok.nl/introductie/-/article/actueel-hoogtebestand-nederland-ahn3- |
| Resolution (m) | | | 1 | 1 |

* CRS = coordinate reference system

*Table S2.2: Predefined vegetation types for the SVM, for each country.*

| | Trees | Shrubs | Heathland | Grassland | Marram | Grey dunes | Bare sand |
|---|---|---|---|---|---|---|---|
| UK | X | | X | X | X | X | X |
| FR | X | X | | X | X | | X |
| BE | X | X | | X | X | X | X |
| NL | X | X | X | X | X | X | X |

*Table S2.3: Overview of the performance parameters of the SVM, per country.*

| Country | Accuracy | 95%-CI | Kappa |
|---|---|---|---|
| UK | 0.9871 | 0.9856, 0.9885 | 0.9846 |
| FR | 0.9842 | 0.9826, 0.9857 | 0.9803 |
| BE | 0.9283 | 0.9241, 0.9324 | 0.914 |
| NL | 0.9314 | 0.9284, 0.9344 | 0.92 |

*Tables S2.4: The confusion matrix performance on the training dataset.*

| UK | | trees | heath | marram | moss | sand | grassland | UA |
|---|---|---|---|---|---|---|---|---|
| | trees | 3830 | 0 | 0 | 1 | 0 | 37 | 0,990176 |
| | heath | 0 | 3988 | 1 | 0 | 0 | 0 | 0,999749 |
| | marram | 0 | 10 | 3993 | 0 | 0 | 21 | 0,992296 |
| | moss | 96 | 0 | 0 | 3993 | 0 | 55 | 0,963562 |
| | sand | 0 | 0 | 0 | 0 | 4000 | 0 | 1 |
| | shrubs | 74 | 2 | 6 | 6 | 0 | 3887 | 0,977862 |
| | PA | 0,9575 | 0,9970 | 0,9983 | 0,9983 | 1 | 0,9718 | |

| FR | | shrubs | trees | marram | sand | grassland | UA |
|---|---|---|---|---|---|---|---|
| | shrubs | 4880 | 126 | 0 | 0 | 0 | 0,974830 |
| | trees | 120 | 4870 | 0 | 0 | 0 | 0,975952 |
| | marram | 0 | 4 | 4933 | 0 | 77 | 0,983845 |
| | sand | 0 | 0 | 0 | 5000 | 0 | 1 |
| | grassland | 0 | 0 | 67 | 0 | 4923 | 0,986573 |
| | PA | 0,9760 | 0,9740 | 0,9866 | 1 | 0,9846 | |

| BE | | shrubs | trees | marram | moss | sand | grassland | UA |
|---|---|---|---|---|---|---|---|---|
| | shrubs | 2031 | 98 | 0 | 7 | 0 | 126 | 0,897878 |
| | trees | 29 | 2269 | 0 | 0 | 0 | 0 | 0,987380 |
| | grassland | 400 | 36 | 0 | 76 | 0 | 2366 | 0,822099 |

| | | | | | | | | | |
|---|---|---|---|---|---|---|---|---|---|
| | marram | 17 | 5 | 2430 | 88 | 0 | 0 | 0,956693 | |
| | moss | 23 | 92 | 70 | 2329 | 0 | 8 | 0,923473 | |
| | sand | 0 | 0 | 0 | 0 | 2500 | 0 | 1 | |
| | PA | 0,8124 | 0,9076 | 0,9720 | 0,9316 | 1 | 0,9464 | | |
| NL | | shrubs | trees | heath | marram | moss | sand | grass | UA |
| | shrubs | 3751 | 22 | 0 | 103 | 103 | 4 | 22 | 0,938924 |
| | trees | 10 | 3977 | 0 | 0 | 0 | 0 | 1 | 0,997242 |
| | grass | 46 | 1 | 0 | 2 | 31 | 0 | 3724 | 0,97897 |
| | heath | 1 | 0 | 3992 | 0 | 0 | 0 | 0 | 0,99975 |
| | marram | 140 | 0 | 7 | 3183 | 321 | 19 | 174 | 0,828044 |
| | moss | 7 | 0 | 1 | 503 | 3477 | 1 | 83 | 0,85388 |
| | sand | 45 | 0 | 0 | 209 | 68 | 3976 | 6 | 0,923792 |
| | PA | 0,9378 | 0,9943 | 0,9980 | 0,7958 | 0,8693 | 0,9940 | 0,9310 | |

S2.3 Quantifying spatial configuration of marram grass

To represent the spatial configuration of marram grass in the study region, two parameters were calculated which are based on an underlying grid. Since the resolution of the aerial photographs differed, four rasters (one per country) were generated with varying grid cells and thus also a varying number of pixels contained within one grid cell (see table S2.1). The corners of the grid cells were chosen as integer numbers within the local Cartesian reference system.

To two calculated landscape metrics were the proportion of marram grass in the area (P) and the spatial autocorrelation or aggregation of marram grass patches in the area (JCS, join-count statistics; see below). The proportion of marram grass in the area was calculated as the ratio between the number of pixels defined as marram grass (A) and the total number of pixels in the raster cell/circle (T).

$$P = A/T$$

The spatial autocorrelation or aggregation was determined based on the join-count statistic (Kabos & Csillag, 2002). This statistic is used with binomial data (such as 1/0 or in this case marram grass/no marram grass) to quantitatively determine the degree of clustering or dispersion of patches. It calculates the sum of joins, thus the number of boundaries of paired raster cells (e.g. pixels) . The possible joins that can occur are: 1-1 , 1-0 and 0-0 joins. For the spatial autocorrelation of marram grass, only the sum of the 1-1 joins is of interest and is calculated as:

$$MM=(1/2) * \Sigma \mathrm{i} \Sigma \mathrm{j} \; (w_{ij} * x_i * x_j)$$

Here $x$ is 1 or 0 when the raster cell is respectively marram grass or not. $wij$ is the spatial weight and is 1 when cell *i* and *j* share a common boundary and 0 otherwise. The z-score is used as a measure for the spatial aggregation of marram grass and is calculated as:

$$Z = (MM_{Oberved} - MM_{Expected}) \; / \; \sigma_{Expected}$$

where the observed sum of 1-1 joins is compared to the expected sum of joins when patches are randomly distributed. Positive z-values correspond to a positive spatial correlation with marram grass patches occurring clustered. Z-values close to 0 correspond to a random distribution of the marram grass patches. Negative z-values correspond to a negative spatial correlation with marram grass patches occurring more regularly distributed than random (Fortin & Dale, 2009).

Both landscape metrics were calculated using R (version 3.6.1, R Core Team 2019). The proportion of marram grass present is calculated using package SDMTools (version 1.1-221.1, VanDerWal 2018). The JCS is calculated by the spdep package (version 1.1-3, Bivand et al. 2018).

The spatial distribution of marram grass in the studied region has been integrated in the following tool: http://www.vliz.be/projects/endure-viewer/. The geographic representation of these statistics is only based on true foredune plots, consisting of sand-marram cover, so omitting already fixed dunes, or those foredunes colonised by (invasive) shrubs.

### *Supplementary material 3: a detailed overview of the simulation model*

*Table S3.1: overview of implemented parameters.*

| **Parameter** | **Definition** | **Unit** | **Default value** |
|---|---|---|---|
| $h$ | Local vegetation height | m | |
| $l_{sat}$ | Saturation length | m | |
| $q$ | Sand flux | kg/m/s | |
| $q_s$ | Saturated sand flux | kg/m/s | |
| $u_*$ | Shear velocity | m/s | |
| $\rho_{veg}$ | Local vegetation density | % | |
| $\tau_s$ | Surface shear stress | Pa | |
| $a$ | Sprouting effect | - | 5 |
| $C$ | Empirical constant to account for the grain size distribution width | - | 1.8 |
| $d_n$ | Nominal grain size | µm | 335 |
| $D_n$ | Reference grain size | µm | 250 |
| $d_{sand}$ | Bulk density of sand | $kg/m^2$ | 1500 |
| $g$ | Gravitational constant | $m/s^2$ | 9.81 |
| $H$ | Maximum vegetation height | m | 0.9 m |
| $s$ | Side length of one cell | m | 0.2 |
| $u_{*th}$ | Shear velocity threshold | m/s | 3.87 * $\alpha$ |
| $u_z$ | Wind speed at height z | m/s | 6.41 |
| $z$ | Height above the bed at which the wind speed is measured | m | 10 |
| $\alpha$ | Conversion factor from free-wind velocity to shear-wind velocity | - | 0.058 |
| $\rho_a$ | Air density | $kg/\ m^3$ | 1.25 |
| $\Gamma$ | Roughness factor of vegetation | - | 16 |

#### *Spatial and temporal dimensions*

The landscape represents a square grid with each cell having a dimension of 0.20 x 0.20 $m^2$. One time step corresponds to one day.

#### *Sand dynamics*

1. Wind direction and boundary conditions

Four different wind directions are defined in the model, each corresponding to a side of the landscape. The distribution of wind direction is assigned at the start of a simulation. Per time step, the wind direction is randomly drawn, based on this distribution. The amount of sand, blown into the system from the sea (N), is expressed as a relative percentage of $q_s$. For instance, if sand influx is defined as 0.5. Then, $q$ equals 0.5 $q_s$ when wind blows from the direction of the sea. Southern winds (land) have an influx of 0 kg sand per cell. Lateral winds have an influx that corresponds with the most recent outflux of a lateral wind, so simulating equal incoming and outcoming lateral fluxes. This amount is constantly updated during a simulation. Wind speed is drawn daily from a normal distribution, based on average wind speed and its standard deviation of that month.

2. Determine shear velocity based on wind velocity (Hoonhout, 2016):

$$u_* = \alpha\, u_z \qquad \text{(eq. 1)}$$

3. Determine maximum (unperturbed) shear stress based on formula for a flatbed (Durán et al., 2010).

$$\tau_* = {u_*}^2 * \rho_a \qquad \text{(eq. 2)}$$

4. Calculate fraction of shear stress acting on the sand, based on density of local vegetation (Duran and Moore, 2013)

$$\tau_S = \frac{\tau_*}{1 + \Gamma\, \rho_{veg}} \qquad \text{(eq. 3)}$$

5. Including Venturi effect: per row of cells occupied by marram grass, perpendicular to the wind direction, the total amount of wind shear stress reduction by the vegetation is calculated. A fraction (25%) of this total amount is then added to the wind shear stress of the two adjacent cells of this row.

6. Recalculate local wind shear velocity based on the formula for an unperturbed shear velocity on a flatbed (Durán et al., 2010):

$$u_* = \sqrt{\frac{\tau_S}{\rho_a}} \qquad \text{(eq. 4)}$$

7. Define saturated sand flux per location based on the Bagnold formula (Bagnold, 1937; Hoonhout, 2016)

$$q_s = C \frac{\rho_a}{g} \sqrt{\frac{d_n}{D_n}} (u_* - u_{*t})^3 \qquad \text{(eq. 5)}$$

8. Erosion is modelled based on the following formula (Kroy et al., 2002):

$$\Delta q_{erosion} = \frac{1}{l_{sat}} q \left(1 - \frac{q}{q_s}\right) \qquad \text{(eq. 6)}$$

In this formula, $l_{sat}$ is assumed to decrease with wind shear according to:

$$l_{sat} = \frac{5}{u_*} \qquad \text{(eq. 7)}$$

The maximum amount of sand that can be eroded, is the amount of sand present in a cell.

9. Deposition is modelled according to:

$$\Delta q_{deposition} = 0.5 * (q - q_s) \qquad \text{(eq. 8)}$$

10. Gravity

Maximum angle of repose is 34° (Durán et al., 2010). Each time step, avalanches are simulated in case this angle is exceeded. Then, the excess amount of sand is displaced to one of the neighbouring cells in the direction with the steepest slope. The maximum angle of repose is set to 34° when vegetation is absent (Durán et al., 2010), but increases with vegetation density. As such, avalanches are less prevalent when plant density is high.

11. Shelter effects

Based on vegetation height and sand availability, average slopes (along the wind direction) are determined within the landscapes. In case a lee slope is steeper than 14° (Kroy et al., 2002), a new imaginary slope of 14° is drawn. The area which is covered by this new slope is sheltered. Within this sheltered area, no erosion is allowed.

Although not used in these simulations, the provided codes allows for:

12. A storm event to occur in the middle of a simulation. A cliff erosion simulation can be included with marram grass and sand destroyed in the first 5 m of the landscape, closest to the sea.

13. Rain events to occur with a chance of 20% (prediction of climate change) from the middle of a simulation onwards.

*Marram grass dynamics*

Seasonality in marram grass:

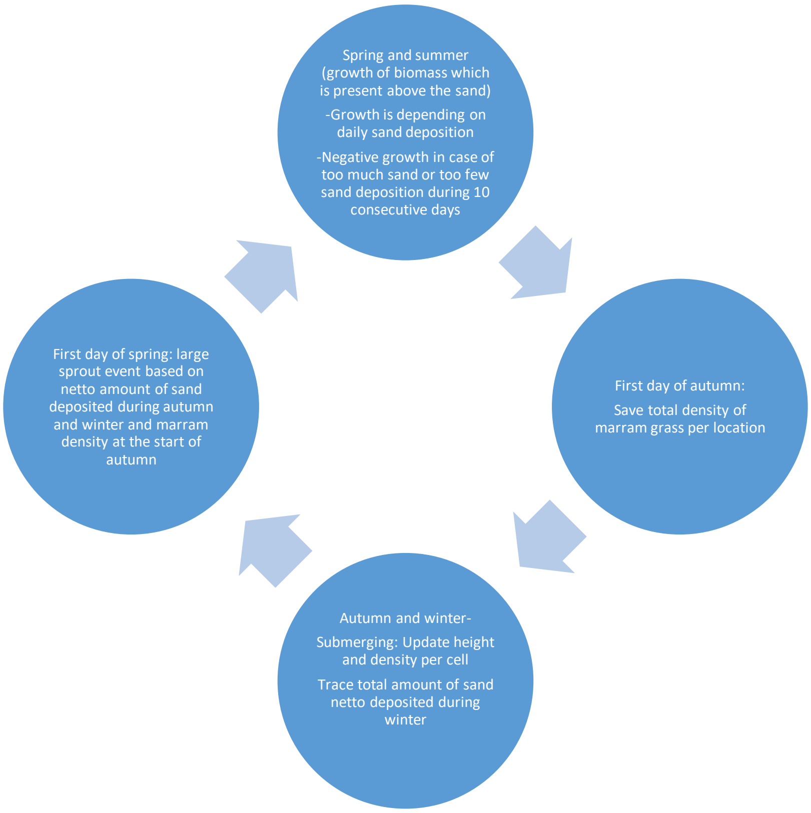

1. Local growth

Marram grass is only able to grow from April to August (for 153 days), according to:

$$\Delta\rho_{veg} = \rho_{veg}\, r\, (1 - \frac{\rho_{veg}}{100}) \qquad \text{(eq. 9)}$$

*r* represents the growth speed of marram grass and depends on the netto amount of sand deposited or removed (by wind and avalanches) during one time step ($\Delta q_{netto}$) (based on (Nolet et al., 2018). In case no netto deposition of sand occurred, growth speed depends on the number of consecutive days without sand deposition ($t_{no\ deposition}$) or too much sand deposition ($t_{too\ much\ deposition}$). To add randomness to the model, an extra value, drawn from a normal distribution with mean 0 and standard deviation 0.01, is added to *r* per cell per time step.

$$r = \left\{ \begin{array}{c} -462.08\left(\Delta q_{netto} - \frac{0.5}{152}\right)^2 + 0.005 \text{ if } \Delta q_{netto} > 0 \\ -0.05 \text{ if } t_{no\ deposition} > 10 \\ -0.05 \text{ if } t_{too\ much\ deposition} > 10 \end{array} \right\} + N(0, 0.001^2) \quad \text{(eq. 10)}$$

2. Lateral growth

During the growth season, marram grass can also grow laterally. The chance of lateral growth ($\gamma$) depends on $\Delta q_{netto}$ according to Nolet et al. (2018):

$$\gamma = -14440\left(\Delta q_{netto} - \frac{0.4}{152}\right)^2 + 0.1 + N(0, 005^2) \quad \text{(eq. 11)}$$

In case lateral growth is successful, one of the eight neighbouring cells is randomly selected as the direction of lateral growth. If marram density in that cell is below 90%, the percentage of marram grass is increased by 1. To add randomness to the model, an extra value, drawn from a normal distribution with mean 0 and standard deviation 0.05, is added to $\gamma$ per cell per time step.

3. Burial from September to March

The height of the vegetation in a cell is estimated by (Van Westen, 2018):

$$h_1 = \sqrt{\rho_{veg}}\ H \quad \text{(eq. 12).}$$

During winter and autumn, vegetation height per cell is updated daily based on the amount of sand deposited or eroded ($\Delta q_{netto}$).

$$h2 = h_1 - \frac{\Delta q_{netto}}{d_{sand}/s^2} \quad \text{(eq. 13).}$$

Afterwards, local vegetation cover is estimated based on $h_2$ by (Van Westen, 2018):

$$\rho_{veg} = (\frac{h_2}{H})^2 \quad \text{(eq. 14).}$$

4. Sprouting event at the start of spring:

The local density of marram grass after sprouting event is determined by the following equation:

$$\rho_{veg,sprout} = \rho_{veg}\,(1 - \Delta h_{winter}) \cdot a \quad \text{(eq. 15).}$$

With $\Delta h$ representing netto change in sand height during autumn and winter. Moreover, in case more than 1 m of sand was locally deposited, marram density becomes 0. In case $\Delta h$ is negative, marram density is unchanged. $a$ determines the strength of the sprouting effect and was set to five based on field observations.

**The code is available on Github:** https://github.com/dbonte/EndureModel.git. This code also includes extra functions, not used here and neither optimized, with regard to erosion dynamics after a simulated storm and germination of marram under wet spring conditions.

Van Westen, B. 2018. Numerical modelling of aeolian coastal landform development.

Van Westen, B. et al. 2019. AEOLIAN MODELLING OF COASTAL LANDFORM DEVELOPMENT. - Coastal Sediments 2019: 1354–1364.

### *Supplementary material 4: Observed changes in dune topography in relation to marram grass spatial organisation.*

The analysis of the evolution of the coastal dunes is based on available high-resolution topographic and morphological data obtained from airborne LiDAR surveys (Strypsteen et al., 2019b) with densities of at least 1 point per m² carried out at annual intervals between 2015 and 2019. In order to scale the maps to those of the maps containing the characteristics of measured vegetation data, each LiDAR survey is converted using linear interpolation into the same grids with a 20x20 m² cell resolution. From these gridded LiDAR surveys elevation differences (*Δheight*) are calculated between the years 2015 and 2019, as well as the coefficient of variation (*CV*) of these differences within each grid cell. The latter thus represents changes in local topography. A subset of the data was taken to fit a linear statistical model: 14 km of coast from the French-Belgian border (De Panne) to the estuary of the Yser (Nieuwpoort). This part of the Belgian coast has the largest dune areas and a minimum of expected disturbance from recreation and beach-cleaning.

To relate *Δheight* and *CV* of the height of the dunes to the cover (*P*) and aggregation of marram grass (join count, *JC*) (see supplement S2), linear models were fitted to the data. A general linear model with a Gaussian distribution was fitted for *Δheight* and a gamma distribution with log-link function for the *CV*. The model was fitted in R v3.5.1 (R Core Team, 2020) using INLA, a Bayesian approach that allows to adjust for spatial autocorrelation while estimating the effect sizes (Rue et al. 2009, Lindgren and Rue 2011, Martins et al. 2013). Fixed effects were the same for the two types of models: linear terms, quadratic terms and interaction terms (excluding the interactions of both quadratic terms) of cover (*P*) and aggregation (*JC*), one linear term for distance to high water mark (*Dist*) and one linear term for mean sand suppletion (*meandV*). The latter are needed to control for regional differences in sand input and or landward ceasing sand fluxes as known sources of potential autocorrelation. Such a control is needed to allow comparison with the simulation model in which processes of sand fluxes are standardised across replicates (simulations thus represent sites directly connected to the beach, experiencing the same sand input). All covariates were standardized before analysis. Priors for the spatial effect, which determine the smoothness of the spatial field, were manually optimized to prevent overfitting (Beguin et al. 2012). This was done because a too flexible spatial effect can adjust for all residual variation which results in a perfectly fitting model with meaningless fixed effects. Predictions were made for combinations of the realised cover and autocorrelation conditions (see main text Fig. 3)

Similar statistical models were fitted to the output of the simulations from the simulation model (see main manuscript and supplement 3). Response variables here are *volume* (comparable to *Δheight* above) and coefficient of variation (*CV*) of volume. Only cover (*P*) and aggregation (*JC*) are used as covariates and no spatial effect was added.

Effect sizes for both the IBM data and LiDAR data are visualised in Fig. S4.1. Spatial fields are plotted in Fig. S4.2, which show the amount of variation that is explained by the spatial effect for different locations, or the extend of the adjustment for spatial correlation at different locations.

Variograms of the residuals for the LiDAR data are shown in Fig. S4.3. Here, the flatter the curve, the less spatial autocorrelation remains after model fitting.

R-script for the statistical models can be found on Github: https://github.com/FemkeBatsleer/DuneTopoMarram

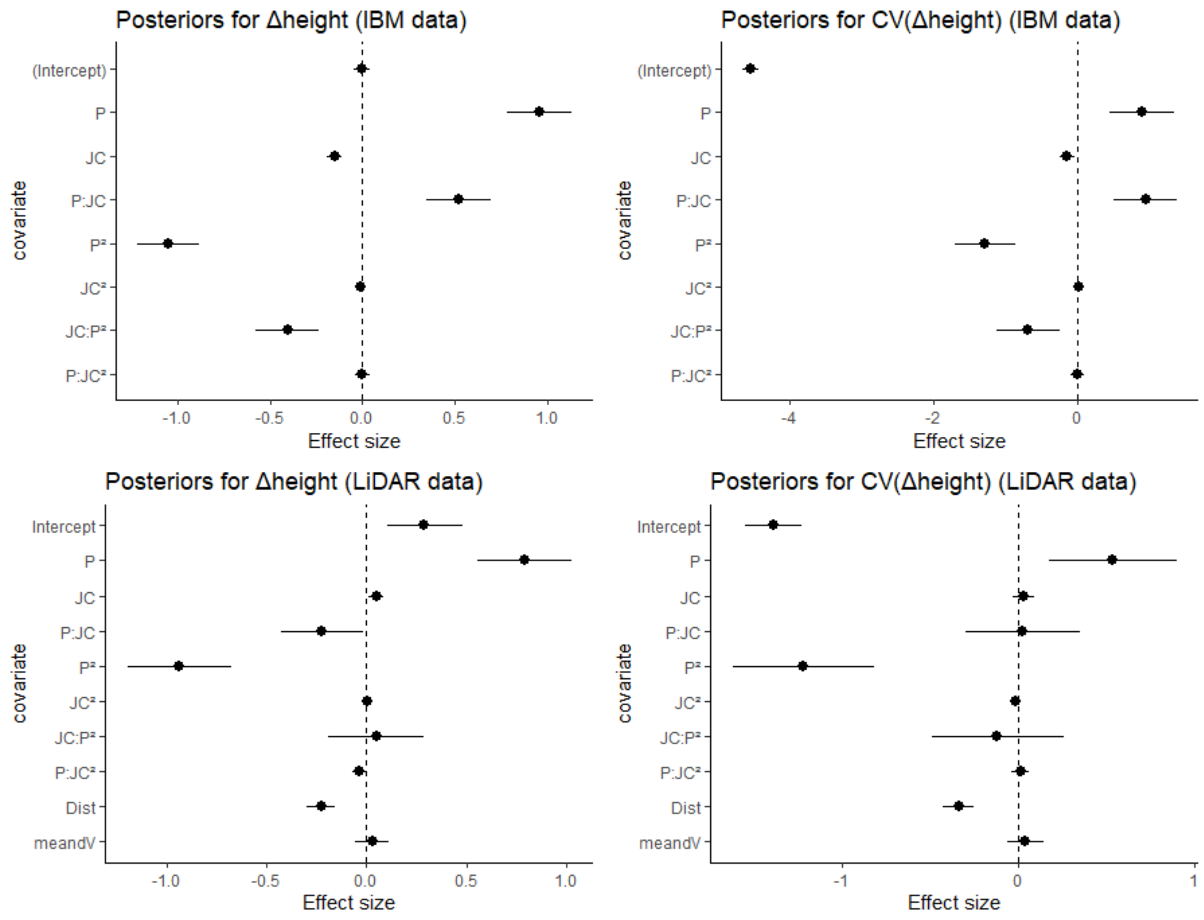

Fig S4.1: Effect sizes of the standardized covariates for the simulations of the IBM model (upper panels) and the LiDAR data (bottom panels. Error bars are credible intervals obtained from posterior estimations with INLA (95% CI: 2.5% quantile and 97.5% quantile). Response variables are *Δheight* (left panels) and coefficient of variance (CV) of the height differences (right panels). Covariates for both IBM and LiDAR data are cover of marram (*P*), clustering of marram (join count, *JC*). The statistical model for the LiDAR data had two extra variables of possible concern in the field study system: distance to the high water mark (*Dist*) and the average of sand suppletion between the two focal years (*meandV*). Note the higher effect sizes for the volume (IBM data) because the response is a volume ($m^3$) and not standardized.

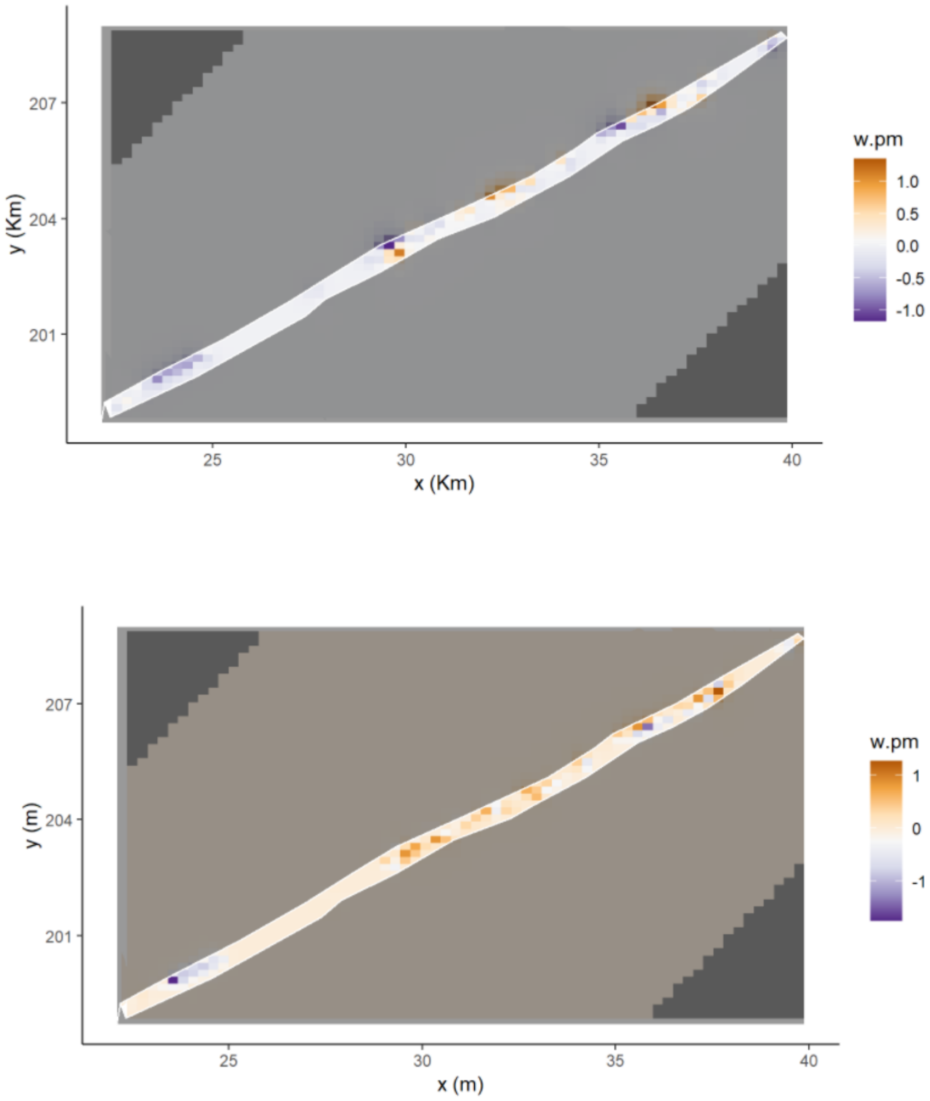

Fig 4.2: spatial field for the LiDAR data models of *Δheight* (top) and *CV* (bottom). This shows spatially how much variance, on top of the fixed effects, is explained by the spatial effect (w.pm). Or in other words, how much the model is adjusted for spatial autocorrelation at those locations.

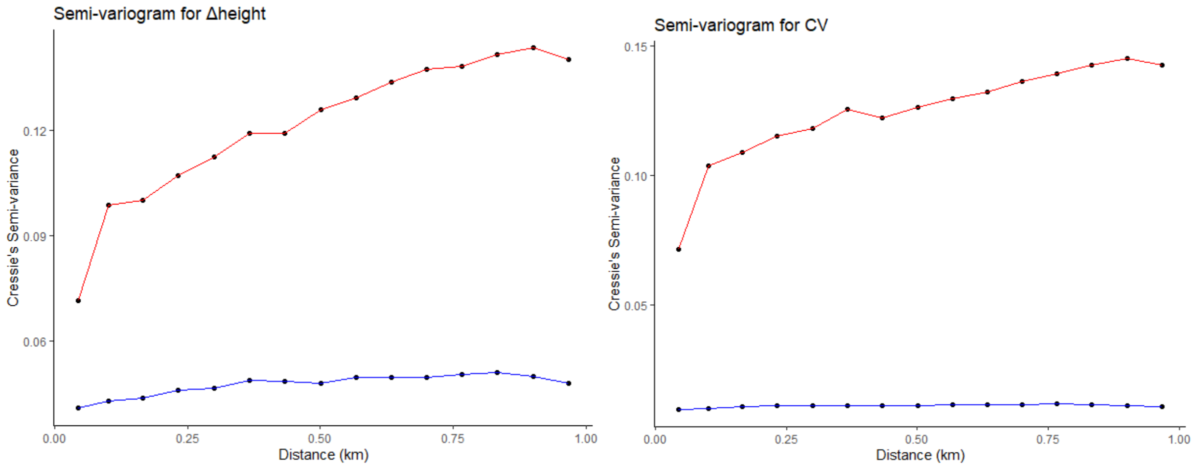

Fig S4.3: Semi-variogram of residuals for non-spatial model (red) and spatial model (blue) for *Δheight* (left) and *CV* (right).

Beguin, J., Martino, S., Rue, H., & Cumming, S. G. (2012). Hierarchical analysis of spatially autocorrelated ecological data using integrated nested Laplace approximation. Methods in Ecology and Evolution, 3(5), 921–929. doi:10.1111/j.2041-210X.2012.00211.x

Lindgren, F., & Rue, H. (2011). An explicit link between Gaussian fields and Gaussian Markov random fields: the stochastic partial differential equation approach. Journal of the Royal Statistical Society B, 73(4), 423–498. doi:10.1111/j.1467-9868.2011.00777.x

Martins, T. G., Simpson, D., Lindgren, F., & Rue, H. (2013). Bayesian computing with INLA: New features. Computational Statistics and Data Analysis, 67, 68–83. doi:10.1016/j.csda.2013.04.014

R Core Team. (2020). R: A Language and Environment for Statistical Computing. R Foundation for Statistical Computing. https://www.r-project.org/. Vienna, Austria: R Foundation for Statistical Computing. Retrieved from https://www.r-project.org/

Rue, H., Martino, S., & Chopin, N. (2009). Approximate Bayesian inference for latent Gaussian models by using integrated nested Laplace approximations. Journal of the Royal Statistical Society B, 71(2), 319–392. doi:10.1111/j.1467-9868.2008.00700.x
